# Durotactic Migration Driven by Anisotropic Matrix Stiffening and Mechanical Feedback

**DOI:** 10.64898/2026.05.19.726229

**Authors:** Donghyun Yim, Brandon Slater, Taeyoon Kim

**Affiliations:** Weldon School of Biomedical Engineering, Purdue University, 206 S. Martin Jischke Dr., West Lafayette, IN 47907, USA

## Abstract

Cell migration is fundamental to various biological processes, including morphogenesis, wound healing, and cancer metastasis. Durotaxis—directed migration of cells in response to spatial variations in stiffness—has been extensively studied using engineered substrates with prescribed stiffness. However, recent work has increasingly shifted toward understanding cell migration in fibrous matrices that can be actively remodeled by the actomyosin contractility, as commonly observed in tumor and epithelial cells. Despite these advances, a theoretical framework explaining how cells structurally remodel their surrounding matrix to promote their own durotaxis, and which cellular forces govern this behavior, remains elusive. To address this gap, we developed a biomechanical model in which polarized cells contract and migrate over a fibrous matrix. Using this model, we first confirmed that cells on an externally strained matrix preferentially migrate along the direction of applied strain. Then, we investigated how cells autonomously remodel the matrix to create stiffness patterns favorable for durotaxis. In the presence of intercellular adhesion, cells acted collectively to stiffen the matrix, after which a small subset of cells escaped the main population and migrated outward. This behavior is reminiscent of intravasation during cancer metastasis, where cohesive cell clusters generate local matrix remodeling that facilitates the departure of more motile subpopulations. These results illustrate how matrix stiffening driven by cell cohesion and contractility regulates durotactic behavior and provide mechanistic insight into collective invasion processes relevant to cancer metastasis.

## INTRODUCTION

Cell migration underlies various biological processes, including wound healing, immune responses, and cancer invasion ^1-3^. Cell migration can be guided by diverse types of cues. Cells preferentially migrate up gradients of ECM stiffness (durotaxis) ^4,5^, chemoattractant concentration (chemotaxis) ^6,7^, or ligand density (haptotaxis) ^8^. Anisotropic structural features of the environment, such as fiber alignments, bias migratory trajectories (contact guidance) ^9,10^. These guidance mechanisms can be additive or competitive. For example, in a matrix consisting of aligned fibers, contact guidance can enable cells to undergo durotaxis in the direction of fiber alignment ^10^, whereas contact guidance and durotactic cues are misaligned, durotaxis tends to predominate in determining migration direction ^11^.

Cell migration is a dynamic, multistep process ^1,12,13^ which begins with cell polarization, the establishment of a spatially asymmetric structure characterized by leading and trailing edges ^1,13^. At the leading edge, localized activation of signaling pathways, including the Rac pathway, drives actin polymerization and the formation of protrusive structures, such as lamellipodia and filopodia ^14^. These protrusions are engaged with an extracellular matrix (ECM) by forming focal adhesions via integrin ^15,16^. Contractile forces generated through the activation of the Rho signaling pathway and Rho-associated kinase (ROCK) are transmitted to the ECM through the focal adhesions ^9,17,18^, evidenced by traction forces measured in experiments ^19^. As a result, the local region of ECM is deformed, resulting in fiber alignment, bundling, and stiffening ^20-22^. Cell-mediated deformation of the ECM has been extensively investigated both in experimental ^23-25^ and computational studies ^26-31^. Experimental studies further demonstrate that cell-induced matrix stiffening can, in turn, generate a mechanically reinforced microenvironment that enhances directional migration by amplifying local mechanosensing cues^9,32^. The metastatic behavior of cancer cells has been extensively investigated in this context. Solid tumor spheroids induce radial alignment and stiffening of the surrounding ECM fibers ^17,18,25,33^, and then cancer cell invasion is enhanced by contact guidance along aligned ECM fibers ^34,35^. By contrast, cell migration is significantly reduced when the matrix is artificially oriented in a tangential direction ^24^.

Several computational approaches have provided important insights into understanding how cells mechanically sense, interact with, and navigate environments ^26-30^. Durotaxis was implemented by allowing cells to form focal adhesions to nearby fiber nodes, calculate local node stiffness, and then migrate toward stiffer nodes ^26,28^. Other models have incorporated fiber alignment-induced contact guidance by updating cell polarity based on the local orientation of neighboring fibers ^27^. A two-dimensional (2D) vertex model was used to capture mechanical interactions between a multicellular system and an underlying elastic network, showing how substrate stiffness regulates collective migration ^36^. Most of the previous models did not rigorously incorporate mechanical interactions between cells and their surrounding environment rigorously; cells were merely prescribed to migrate toward stiffer regions or along aligned fibers. To address these limitations, we previously developed a mechanistic, physics-based model ^37^. In the model, a single polarized cell generates protrusions in random directions on a viscoelastic fibrous matrix and applies contractile forces to matrix nodes located beneath protrusions. Matrix nodes with higher elasticity resist the applied contractile forces more strongly, thereby promoting migration toward stiffer regions. We used matrices with either soft and stiff regions or stiffness gradient to recapitulate durotactic migration patterns observed experimentally and define the mechanism of durotaxis.

To further elucidate how cell migration emerges from dynamic cell–matrix feedback, we extended our previous model to examine two complementary scenarios. First, we examined migratory patterns of multiple cells on an externally strained matrix with anisotropic stiffness to understand why cells preferentially migrate in stiffer directions. Then, we investigated how cells remodel and anisotropically stiffen their surrounding matrix by exerting contractile forces.

## MATERIALS AND METHODS

### Simplification of cells and an underlying matrix

A migrating cell is represented by a polarized unit consisting of front and rear cell-points (Fig. 1A). A cell polarity vector was defined from the rear cell-point to the front one. Each cell-point has a semicircular (donut half-shaped) adhesion zone defined by outer (*R*_out_) and inner (*R*_in_) radii. To capture the asymmetric size of polarized cells, the outer and inner radii of the adhesion zone for the front cell-point are assumed to be larger than those for the rear cell-point, such that

**Figure 1.**
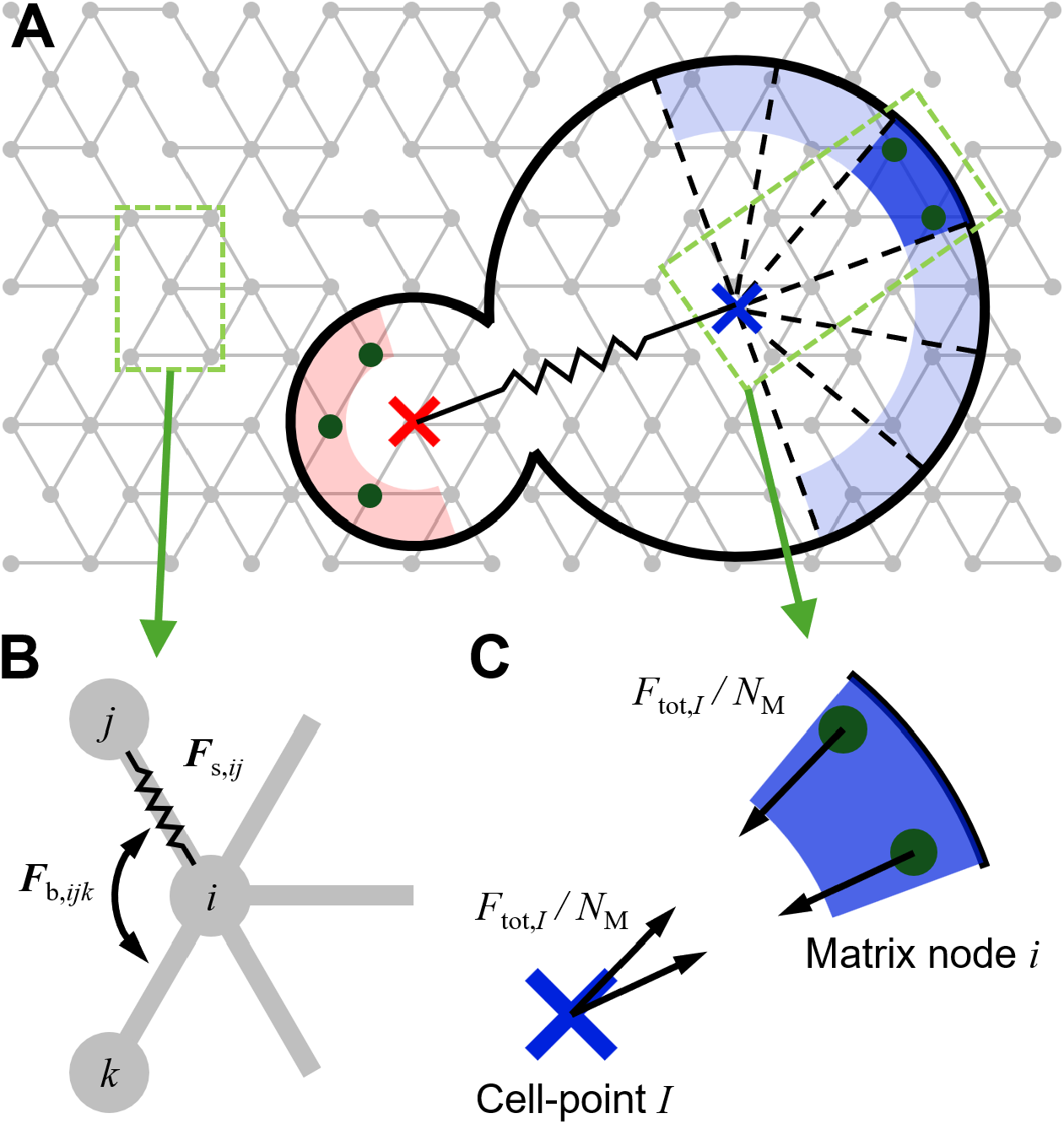
Cell migration model. (A) Each cell is simplified into a unit with front and rear cell-points whose inter-distance is maintained by a spring force. Cell polarity is defined by a vector from a rear cell-point to a front one. Each cell-point has an annular adhesion zone characterized by outer and inner radii and the angle of 180°. The adhesion region of the front cell-point is divided into 6 sub-regions, each of which represents a lamellipodium. One of the 6 sub-regions is randomly selected for activation, and then the activated sub-region lasts for 0.1 h before switching to another sub-region for activation. (B) An underlying matrix is discretized into nodes initially placed on a triangular lattice and interconnected by chains. The polymeric and plastic natures of an extracellular matrix are considered by randomly removing a portion of the chains. The elastic aspect of the matrix is governed by extensional (*F*_s,*ij*_) and bending force (*F*_b,*ijk*_) acting on the matrix chains. (C) Each cell-point contracts all matrix nodes present within an activated sub-region by applying a contractile force (*F*_c,*iI*_) which is equal to the total contractile force (*F*_tot,*I*_) divided by the number of interacting matrix nodes (*N*_M_). Contraction of the front cell-point is stronger than that of the rear cell-point to account for stronger contraction near the leading edge than the trailing edge (i.e., *F*_tot,F_ > *F*_tot,R_). Each cell-point experiences a reaction force from interacting matrix nodes (***F***_c,*Ii*_ = −***F***_c,*iI*_), meaning that the cell point is pulled toward the matrix nodes.

*R*_out,F_ > *R*_out,R_ and *R*_in.F_ > *R*_in,R_. An underlying matrix is modeled as a network of nodes initially arranged on a triangular lattice and connected by chains. The polymeric nature of ECM is incorporated by randomly removing a portion of chains with given probability (*p*) ^38^. Once established, matrix connectivity is assumed to remain fixed over time.

### Model formulation

In our model, we assume an overdamped system in which inertial contributions are negligible compared to viscous drag over time scales considered in our simulations. Viscoelastic interactions are represented using the Kelvin–Voigt element, consisting of a linear spring and a dashpot arranged in parallel. All parameters and their values used in simulations are listed in Table S1.

Each governing equation for cell-points and matrix points has two types of forces:

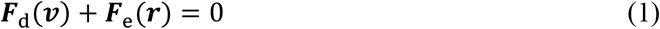

where ***F***_d_ represents viscous drag forces depending on velocities, ***v***, and ***F***_e_ represents elastic forces determined by positions, ***r*.**

The total number of the governing equations is 2(*N*_CP_ + *N*_M_), where *N*_CP_ and *N*_M_ represent the numbers of cell-points and matrix points, respectively, and the coefficient 2 originates from the use of two-dimensional space. Collectively, all the governing equations can be reduced into a matrix equation form comprising linear equations:

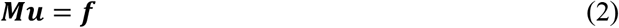

where ***M*** is a matrix with drag coefficients involved with drag forces, and ***u*** is an unknown vector containing the x and y components of velocities of cell-points and matrix points, and ***f*** is a vector consisting of the x and y components of velocity-independent forces (Fig. S1). By solving the matrix equation, the velocities of the cell-points and matrix points are obtained at each time step, and then their positions are updated using the velocities and the Adams-Bashforth fourth-order integration scheme ^39^.

### Governing equation and forces for cell-points

For each cell-point *I*, force balance is considered by the following governing equation:

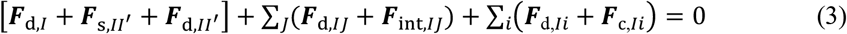

where *J* indicates other cell-points adjacent to *I*, and *i* indicates matrix points interacting with *I*. The first 3 terms within brackets are forces acting on each cell without interaction with other cells. Two forces in the first summation notation (Σ_*J*_) are forces stemming from interactions with neighboring cells. The last terms within the other summation notation (Σ_*i*_) are forces originating from interactions with a matrix.

***F***_d,*I*_ is a drag force generated by a surrounding viscous medium, defined as:

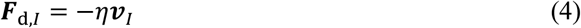

where *η* is a cell drag coefficient, and ***v***_*I*_ is the velocity of a cell-point *I*. ***F***_s,*II*′_ is a spring force maintaining a distance between front and rear cell points (= *r*_*II*′_) close to its equilibrium value (*r*_0,FR_) with extensional/compressive stiffness (*κ*_s,FR_), which is derived from the following harmonic potential (*U*_s,*II*′_):

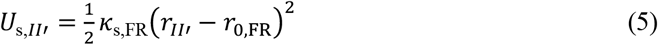

***F***_d,*II*′_ is a viscous force originating from a relative movement between the front and rear cell-points of the same cell (i.e., intracellular drag):

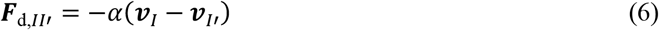

where *α* is a front-rear drag coefficient.

***F***_d,*IJ*_ is a drag force acting between a cell-point *I* and other cell-point *J* that belongs to different cells (i.e., intercellular drag). This drag force acts only when a distance between two cell-points (*r*_*IJ*_) falls below a critical threshold (*r*_cr,int_ = *R*_out,*I*_ + *R*_out,*J*_):

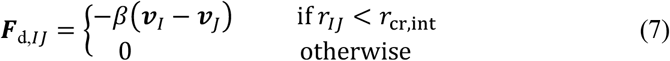

where *β* is a cell-cell drag coefficient. ***F***_int,*IJ*_ is an interaction force which accounts for repulsion (volume exclusion) or attraction (cell-cell adhesion) depending on *r*_*IJ*_ (Fig. S2) ^40^:

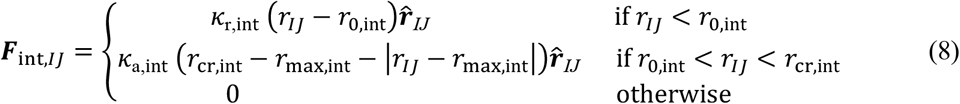

where *κ*_r,int_ and *κ*_a,int_ are the strength of repulsion and attraction, respectively, *r*_0,int_ is an equilibrium distance where neither of them is applied (= *R*_in,*I*_ + *R*_in,*J*_), *r*_max,int_ is a cutoff distance beyond which the attractive force stops acting (= (*r*_0,int_ + *r*_cr,int_)/2), and 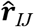 is a unit vector defined from a cell-point *I* to a cell-point *J*. ***F***_d,*IJ*_ and ***F***_int,*IJ*_ are calculated between the cell-point

*I* and all neighboring cell-points *J*.

***F***_d,*Ii*_ is a viscous force acting between a cell-point *I* and a matrix point *i*:

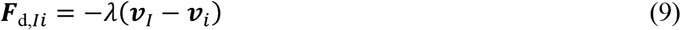

where *λ* is a cell-matrix drag coefficient. This drag force represents the viscous effect of focal adhesion connection, following the prior study that has shown that multiple transient focal adhesion connections can be effectively modeled as frictional forces ^41^. ***F***_c,*Ii*_ is a contractile force that the cell-point *I* exerts to nearby matrix nodes, which will be explained in detail later. ***F***_d,*Ii*_ and ***F***_c,*Ii*_ are calculated for all matrix points interacting with a cell-point *I*.

### Governing equation and forces for a matrix

For each matrix point *i*, force balance is considered by the following governing equation:

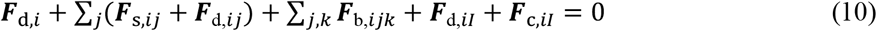

where *j* and *k* indicate other matrix points connected to *i* (Fig. 1B), and *I* indicates a cell-point interacting with *i*. The first 4 terms are forces regulating the elastic or viscous behavior of the matrix. ***F***_d,*i*_ is a drag force generated by a surrounding viscous medium, defined as:

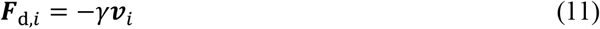

where *γ* is a matrix drag coefficient, and ***v***_*i*_ is the velocity of a matrix point *i*. ***F***_s,*ij*_ is a linear spring force maintaining a distance between two matrix points *i* and *j* (= *r*_*ij*_) close to its equilibrium length (*r*_0,M_) with stiffness *κ*_s,M_, which is derived from the following harmonic potential (*U*_s,M_):

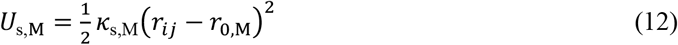

To account for the buckling of matrix fibers under compression ^42^, the nonlinear behavior of matrix fibers is incorporated, as in previous models ^26,30,43^, by assigning asymmetric values to *k*_s,M_:

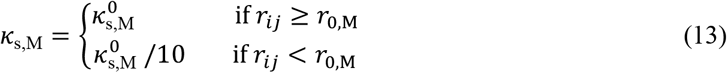

***F***_d,*ij*_ is a drag force acting between matrix points *i* and *j* to account for the viscous aspect of the matrix:

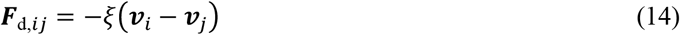

where *ξ* is a matrix-matrix drag coefficient. ***F***_s,*ij*_ and ***F***_d,*ij*_ are calculated between the matrix point *i* and all matrix points connected to *i*.

***F***_b,*ijk*_ is a bending force enforcing an angle formed by three matrix points (*θ*_*ijk*_) to remain close to its equilibrium value (*θ*_0,M_) with bending stiffness (*κ*_b,M_), which is derived from a harmonic potential (*U*_b,*ijk*_):

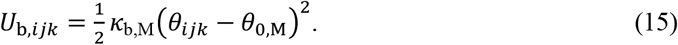

The last 2 terms are involved with interactions with a single cell-point. ***F***_d,*iI*_ and ***F***_c,*iI*_ are reaction forces to ***F***_d,*Ii*_ and ***F***_c,*Ii*_, respectively, meaning their magnitudes are identical, whereas their signs are opposite to each other.

### Mechanical interactions between cells and a matrix

To probe their local environment, cells repeatedly generate lamellipodia-like protrusions in stochastic directions at the front cell-point. The front adhesion region is partitioned into 6 sub-regions, each of which is assumed to correspond to a lamellipodium. One lamellipodium is randomly selected for activation and remains active for 0.1 h before retracting, after which another lamellipodium is activated. By contrast, the entire rear adhesion region is assumed to always remain active.

To represent coupling mediated by focal adhesions, each cell-point applies contractile forces to all matrix nodes located within its activated lamellipodial region (Fig. 1C). Each matrix node is assigned to only one cell-point; if a matrix node lies within the active lamellipodial regions of multiple cell-points, it interacts exclusively with the nearest one. For a given cell point *I* and every matrix point *i* interacting with *I*, contraction exerted by *I* onto *i* is defined by the following equation:

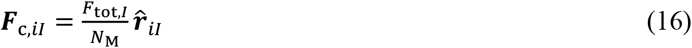

where *F*_tot,*I*_ is a total contractile force (= *F*_tot,F_ for front cell-points and *F*_tot,R_ for rear cell-points), *N*_M_ is the number of matrix points interacting with *I*, and 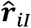 is a unit vector defined from *i* to *I*. Note that the total contraction force of front cell-points (*F*_tot,F_) is much higher than that of rear cell-points (*F*_tot,R_) to drive cell migration forward. Partitioning of the adhesion region for the front cell-point makes the contractile force act on fewer matrix nodes in random directions. As mentioned earlier, each cell-point experiences the corresponding reaction force (***F***_c,*Ii*_) exerted by the matrix nodes, and these reaction forces collectively drive the displacement of the cell-points:

### Simulations with externally strained rectangular matrices

We probe how cell migration responds to anisotropic stiffening of a matrix induced by external strain. We apply normal strain (*ε*) at the left and right boundaries of the rectangular matrix, ranging from 0 to 2.5. The initial dimension of the matrix was set for the final dimension to almost become 1000 µm × 1000 µm after strain application. The number of matrix nodes is set at similar level regardless of *ε* by adjusting the equilibrium chain length (*r*_0,M_). For example, for *ε* = 0, the initial dimension is 1000 µm × 1000 µm, and *r*_0,M_ is 5 µm with 46,431 nodes. For *ε* = 2.5, the initial dimension is 257 µm × 1000 µm, and *r*_0,M_ is 2.67 µm with 46,764 nodes.

Anisotropic stiffening of the matrix is evaluated as follows. Once a matrix is fully relaxed following strain application, we impose external forces (*F*_ext_) on all matrix nodes within a 100 µm × 100 µm square region centered in the network for 1 h in either parallel or perpendicular direction to the strain direction (Figs. S3A, B). Cells are not included for this measurement. The magnitude of *F*_ext_ is set to 0.01 nN, which is similar to force level that each matrix node experiences in simulations for cell migration (i.e., ∼|***F***_c,*iI*_|). Considering stiffness is inversely correlated with displacement under a constant applied force ^44^, the ratio of stiffnesses measured in parallel (*E*_||_) or perpendicular (*E*_*⊥*_) direction to the applies strain is defined as follows:

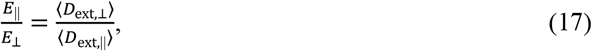

where ⟨*D*_ext,||_⟩ and ⟨*D*_ext,*⊥*_⟩ are the ensemble average of the displacement of matrix nodes subject to *F*_ext_.

At the beginning of simulation, 100 cells are positioned near the center of the matrix within a circular region of radius 50 µm. To achieve relatively uniform initial spacing, the rear cell-point of each cell is assigned to rectangular grids with a spacing of 17.7 µm in both x and y directions. Then, the front cell-point is placed at a distance *r*_0,FR_ from the rear cell-point with a randomly assigned orientation. Simulations are run for a total duration of 100 h. To evaluate the tendency of cells to migrate toward stiffer regions, we employ the durotactic index ^44^:

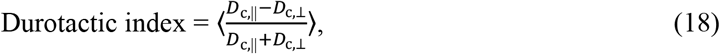

where *D*_c,||_ and *D*_c,*⊥*_denote distances traveled in parallel and perpendicular, respectively, to the direction of *ε* during the simulation. To minimize boundary effects, *D*_c,||_ and *D*_c,*⊥*_ are calculated when cells cross a circular line (radius 450 µm, centered at the domain center) by comparing their initial positions with positions at which the cells cross the circular line. For cells that do not reach the circular line before the end of simulations, *D*_c,||_ and *D*_c,*⊥*_ are calculated by comparing their initial positions with their last positions at the end of simulations.

### Simulations with circular matrices without external strain

To examine migration directionality in the absence of externally applied strain, we model a circular matrix of radius 500 µm. The initial spacing between matrix nodes (*r*_0,M_) is set to 5 µm, and all nodes located on the circular boundary are fixed in space. To avoid boundary effects originating from the non-periodic outer boundary, cells approaching the boundary within a distance of 4*R*_out,F_ are removed from simulations. All other model parameters are identical to those used in the rectangular-domain simulations.

To assess matrix contraction induced by cells, we first define a circular region with radius *R*_a_, set to 200 µm by default. All matrix nodes whose distances from the domain center lie between (*R*_a_ - 1 µm) and (*R*_a_ + 1 µm) are identified, and these nodes are traced throughout the simulation. Matrix strain is then calculated as the mean radial displacement of these traced nodes normalized by *R*_a_. To evaluate force development on the matrix, after matrix chains crossing a circle with a radius *R*_a_ are identified, the magnitude of tensile forces acting on chains which cross a circle with radius *R*_a_ (*F*) is calculated, and an angle between a radial direction and the orientation of the chains (0° ≤ *θ* ≤ 90°) is also computed. Then, the radial (*F*_*r*_) and circumferential (*F*_*θ*_) components of the tensile force are calculated as:

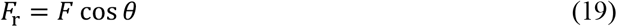

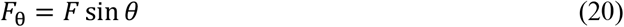

We quantify the fraction of escaped cells by counting the number of cells that remain within a circle of radius *R*_a_ at each time point. Finally, the radial migration bias is computed from the trajectories of cells from a time point when the cells cross the same circle of radius *R*_a_ and a last time point when the migration is ended. Using cell positions with 10-h time delay, the radial migration bias is calculated as:

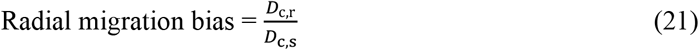

where *D*_c,r_ and *D*_c,s_ are the radial displacement and the shortest distance between cell positions at two different time points (with the 10-h time delay), respectively ^45^.

## RESULTS

### Uniaxial loading induced anisotropic stiffnesses in matrices

We first sought to induce anisotropic stiffening in the initially relaxed substrate by applying strain to both boundaries of the rectangular matrix in one direction, following experimental approaches ^46^ (Fig. 2). The strain was linearly increased and held at a constant level, *ε*. We varied *ε* and the probability of matrix chain removal (*p*) to understand how these parameters affect anisotropic stiffening and plastic deformation (Fig. 2A). With *ε* = 2, matrix chains became horizontally aligned and exhibited greater stiffness in the horizontal than in the vertical direction. When the matrix was fully connected (*p* = 0), all horizontal chains stretched to similar extents. By contrast, when fewer chains were present (*p* = 0.5), a subset of chains bore disproportionately high tension. This occurred because increased stochasticity in connectivity generated chains with fewer parallel load-bearing paths, causing these chains to stretch more—analogous to a compliant spring elongating more than a stiffer one when two springs are connected in series and subjected to the same load. At the same time, the low connectivity limited the number of available tension-transmitting paths, leaving many chains largely relaxed. Conversely, with *p* = 0, the high connectivity allowed tension to be distributed across many horizontal chains, reducing the need for any individual chain to undergo excessive stretching.

**Figure 2.**
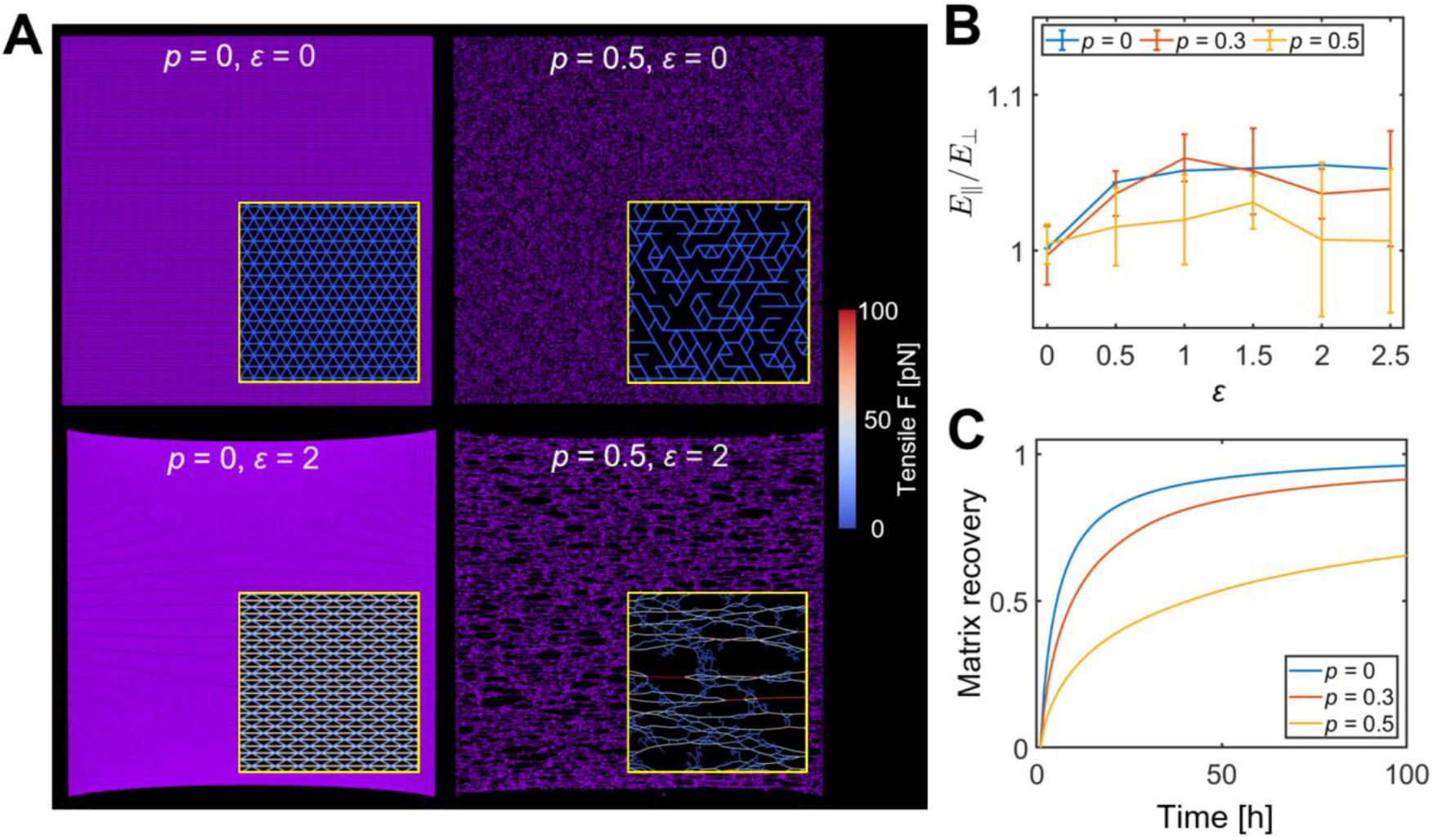
Anisotropic matrix stiffening is induced by external strain. Normal strain is applied to the left and right boundaries of the matrix whose chains were removed at a given probability (*p*), up to a specified strain level (*ε*). Top and bottom boundaries were allowed to freely move. Cells were not included in these simulations. (A) Examples of matrices with different values of *p* and *ε*. With high *ε*, the matrices slightly shrank in the other direction, implying a positive Poisson’s ratio. Insets: magnified views with the magnitude of spring forces visualized via color scaling. When *ε* > 0, some of the matrix chains were highly stretched and aligned along the direction of applied strain. (B) The extent of stiffness anisotropy depending on *p* and *ε*. Matrix stiffness was measured in parallel (*E*_||_) or perpendicular (*E*_*⊥*_) direction to externally applied strain by applying forces to a small portion of matrix nodes and evaluating their displacements. In general, *E*_||_/*E*_*⊥*_ was greater with higher *ε* and lower *p*. (C) Plasticity of matrices depending on *p* and *ε*. Level of matrix recovery was probed with *ε* = 2 by applying forces to a small subset of matrix nodes, releasing the forces, and then comparing their positions with initial positions over time. Plasticity was greater (i.e., lower matrix recovery) at higher *p*.

To quantify stiffness anisotropy (*E*_||_/*E*_*⊥*_) across different values of *p* and *ε*, we applied *F*_ext_ to selected matrix nodes and measured their displacements (Figs. S3A, B). The ratio *E*_||_/*E*_*⊥*_ was then evaluated for different bending stiffness of the matrix (*κ*_b,M_ = 0, 0.001, and 0.01 nN⋅µm) (Figs. 2B and S3C, D). At higher *p*, only a subset of chains stiffened, so the anisotropy signal was diluted when averaged over the entire network; many relaxed chains displaced similarly regardless of the direction of *F*_ext_. Moreover, with intermediate *κ*_b,M_ (= 0.001 nN⋅µm), the peak *E*_||_/*E*_*⊥*_ was greater than that with *κ*_b,M_ = 0.01 nN⋅µm, reflecting that anisotropic stiffening is more effectively induced when the matrix is more compliant to bending deformation. Note that boundaries in the other direction were free to move during these measurements, so the matrix showed a slight decrease in its dimension in the other direction (e.g., *p* = 0, *ε* = 2), indicative of a positive Poisson’s ratio. To check whether these free boundaries could potentially introduce artifacts into the stiffness analysis, we repeated simulations with those boundaries fixed in space (Fig. S3E). The results remained largely consistent, indicating that the influence of the free boundaries on our stiffness-anisotropy measurements was minimal.

The ECM is inherently plastic, meaning that its deformation is not fully reversible once external stress or cell-generated contractility ^47,48^. To evaluate ECM plasticity within our computational framework, we first applied *F*_ext_ and then removed it, allowing the matrix to relax for 100 h across different values of *p* (Fig. 2C). We found that substrate nodes returned to their original positions more slowly when a larger fraction of chains was removed. In these sparsely connected networks, relaxed chains received insufficient restorative forces from neighboring chains to fully recover their initial configuration (e.g., *p* = 0.5, *ε* = 2). To ensure that our simulations captured both stiffness anisotropy and ECM plasticity, we selected intermediate connectivity of *p* = 0.3 for all subsequent analyses unless otherwise noted.

### Stiffness anisotropy in strained matrix guided cell migration

It has been experimentally demonstrated that strained matrices can induce directed cell migration ^44,46,49^. We next examined how stiffness anisotropy influences migration trajectories in our model. Following strain application, cells migrated preferentially along the direction of *ε* (Figs. 3A, B and Movies S1, S2). Quantification using the durotactic index confirmed this trend, showing that cells consistently migrated along the strain direction (Fig. 3C). The durotactic index increased with *ε*, whereas higher values of *p* reduced migration directionality—both trends mirroring the behavior of stiffness anisotropy (Figs. 2B and 3C). At *p* = 0.5, the durotactic index was lower than at smaller *p* values because many cells interacted with relaxed chains in which stiffness anisotropy was absent or only weakly developed.

**Figure 3.**
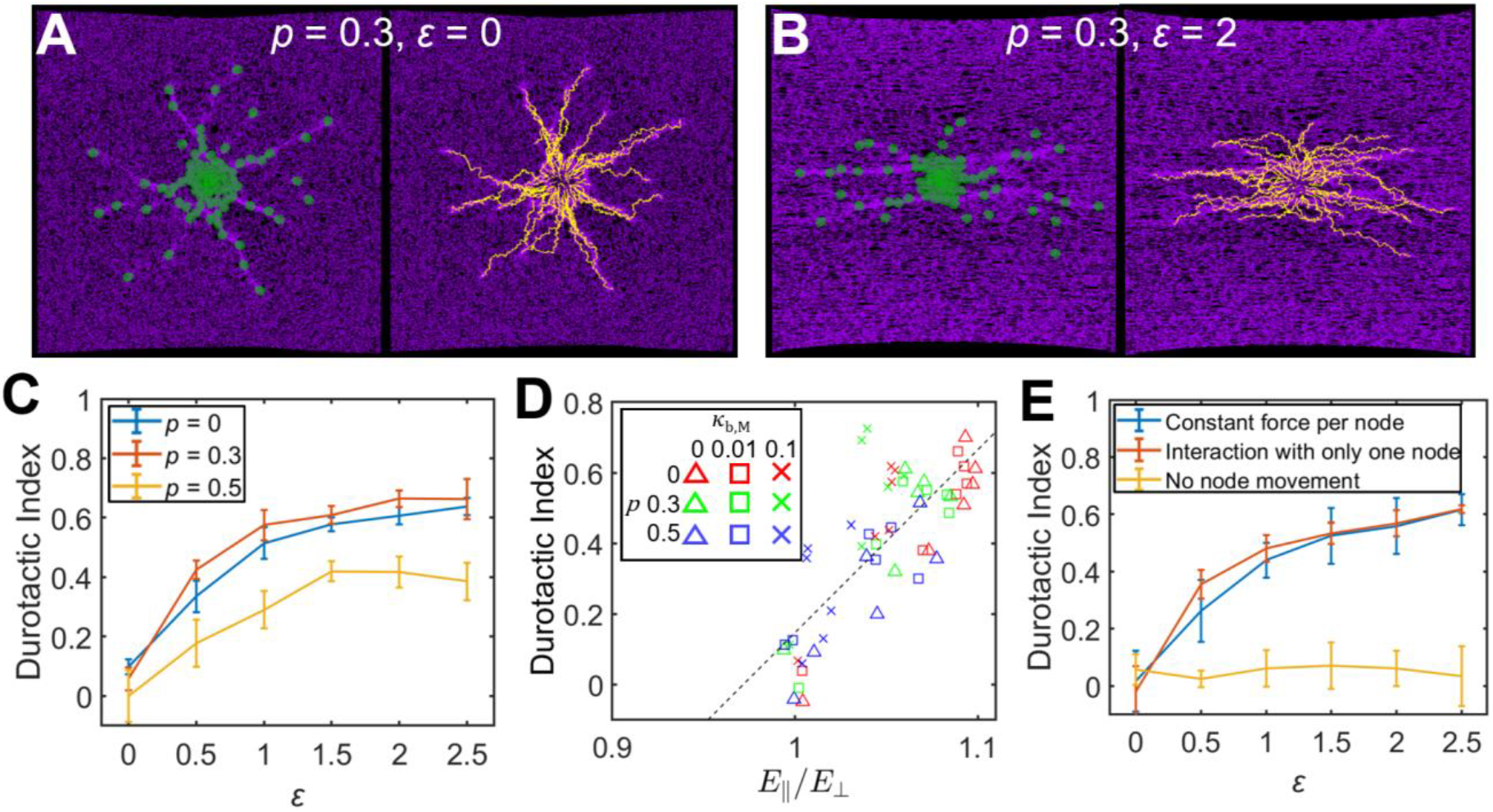
Durotactic migration is mediated by anisotropic stiffening. Cells were allocated at the center of externally strained matrices with various values of *p* and *ε*, and their durotactic migration was evaluated using the durotactic index which ranges between -1 (worst) and 1 (best). (A, B) Examples of cells (left) and their cumulative trajectories (right) were visualized on matrices. (C) Durotactic index as a function of *p* and *ε*. Cells experienced more directed migration along the direction of applied strain with lower *p* and higher *ε*. (D) A correlation between stiffness anisotropy (*E*_||_/*E*_*⊥*_) and the durotactic index. (E) Durotactic index measured using different types of cell-matrix interactions with *p* = 0.3 and various *ε*. Cells did not exhibit durotactic migration when matrix nodes were not allowed to move by forces exerted from the cells.

We repeated the durotactic-index analysis for *κ*_b,M_ = 0 and 0.001 nN⋅µm (Fig. S4). To further clarify a correlation between stiffness anisotropy and directed migration across multiple *κ*_b,M_ and *p* values, we plotted *E*_||_/*E*_*⊥*_ ratios and durotactic indices in all cases (Fig. 3D). Linear regression across all data revealed a positive correlation between these two metrics.

In the presence of stiffness anisotropy, migration paths emerged as leader cells contracted and moved through specific regions of the matrix (Fig. 3B and Movie S2). Follower cells subsequently sensed the locally stiffened paths, enabling coordinated migration behind leader cells. This behavior is consistent with previous computational work showing that one cell can follow the trajectory of another that initiated migration earlier ^27^.

Finally, we tested the robustness of our findings by modifying the mechanism of force transmission between cells and the substrate (Figs. 3E and S5). First, we evaluated an alternative formulation of cell–matrix interaction originally defined in Eq. 16 (Fig. 1C) by applying *F*_c,*iI*_ to all engaged matrix nodes without dividing (Fig. S5A) or to a single matrix node (Fig. S5B). These modifications did not substantially alter the results relative to the original formulation (Fig. 3E). It means that the denser matrix nodes induced by fiber alignment are not the main mechanism for biased migration (i.e., not contact guidance-driven migration). In addition, our model reproduced durotaxis through Eq. 9, in which ***v***_*i*_ reflects the stiffness sensed at substrate point *i* via its interactions with connected cell-points. To eliminate this stiffness-dependent effect on cell-point velocity, we next fixed ***v***_*i*_ = 0 at all times (Fig. S5C). Under these conditions, the durotactic index dropped nearly to zero even after straining, demonstrating that durotaxis is the primary driver of directed migration in our model (Fig. 3E).

### Cell-cell adhesion is required for matrix remodeling and sporadic cell migration

Biological cells typically contract and remodel their surrounding matrix, generating stiffness cues that enable durotaxis. Consistent with this, previous experimental studies of contractile tissues in isotropic matrices have shown that radial matrix alignment and stiffening precede the onset of cell migration ^17,18,33^. In tumor spheroid models using mouse colon carcinoma cells, collagen contraction began immediately after spheroid embedding, whereas outward cell migration occurred only after a delay ^32^. Similarly, breast cancer spheroids were shown to radially align collagen fibers up to five tumor radii away before invasion ^50^. This phenomenon is not restricted to cancer cells: mammary epithelial tissues and fibroblast aggregates also align surrounding collagen prior to migration ^17^.

To test whether our model reproduces this temporal precedence of matrix deformation, we simulated cell migration within a circular domain to eliminate geometric asymmetry while keeping all other parameters identical to the rectangular-domain simulations (Fig. 4A, left labeled with “w/o adh, w/ *F*_tot,F_”). Cells migrated through an initially isotropic matrix, and matrix deformation arose solely from cell-generated contractility (Movie S3). By tracking matrix nodes initially located on a circle with *R*_a_ = 200 µm, we found that inward matrix contraction peaked between 100 h and 200 h and subsequently relaxed toward the original configuration (Figs. 4B, C). Correspondingly, forces developed in radial (*F*_r_) and circumferential (*F*_θ_) directions peaked early and then decayed to zero, with *F*_r_/*F*_θ_ maintained around 2 (Figs. 4D and S6A). This relaxation occurred because, as more cells escaped from the cluster, the matrix recovered its initial configuration through the restoring contributions of extensional and bending forces (Fig. 4E). It was observed that 73% of cells migrated with the radial migration bias greater than 0.8 (Fig. 4F).

**Figure 4.**
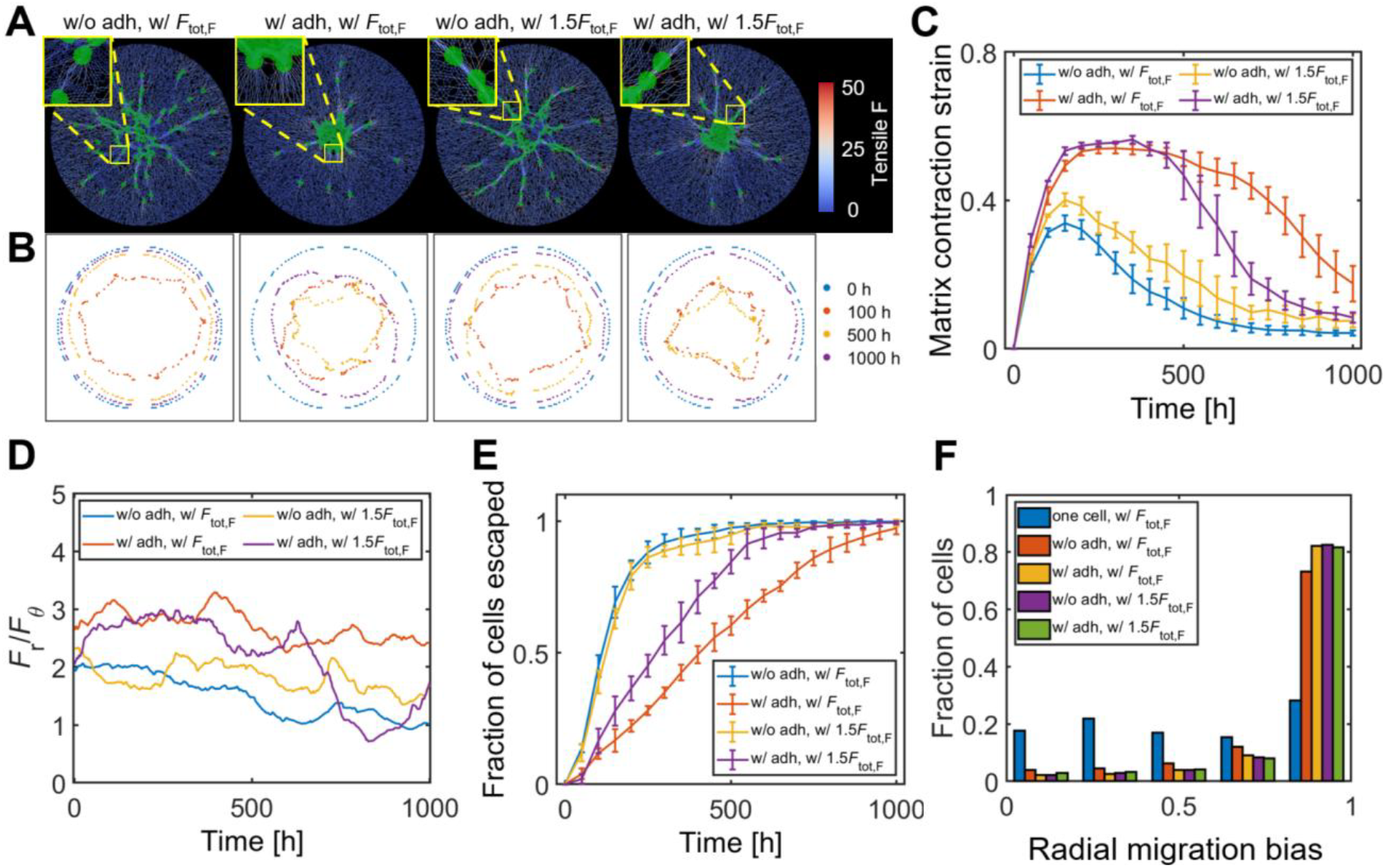
Anisotropic stiffening and radial migration are enhanced by intercellular adhesion and cellular contractility. (A) Cell migration on a circular matrix under four conditions: i) w/o cell-cell adhesion (adh), w/ the reference total contractile force of the front cell-point (*F*_tot,F_), ii) w/ adh, w/ *F*_tot,F_, iii) w/o adh, w/ 1.5*F*_tot,F_, and iv) w/ adh, w/ 1.5*F*_tot,F_. Cells were initially seeded as circular population near the center of the matrix. Cases with cell-cell adhesion exhibited delayed cell escape from the cluster after showing more significant matrix contraction. With higher *F*_tot,F_, cells escaped the cluster more frequently and followed a leader cell (if any) through aligned matrix chains. Snapshots were taken at 100 h. (B) Matrix nodes initially located at 200 µm from the matrix center were traced during simulations under the same four conditions as those in (A). Contraction was more sustained in the presence of cell-cell adhesion. (C) Matrix strain calculated using the traced matrix nodes shown in (B). (D) The radio of radial forces (*F*_r_) to circumferential ones (*F*_θ_) acting on the matrix. (E) Fraction of cells escaping the cell cluster. (F) Radial migration bias indicating the extent of radial migration.

In contrast, when the same analysis was performed for the single-cell case, only 28% of cells had the radial migration bias greater than 0.8 (Fig. 4F, labeled with “one cell, w/ *F*_tot,F_”), suggesting the collective force generation may be required to enhance the radial migration. These results prompted us to ask which mechanistic factors further enable cells to migrate within a stiffened matrix and thereby exhibit durotaxis.

Experimental studies have implicated intercellular adhesion as a key regulator of matrix alignment and collective force transmission. Radial fiber alignment increased when E-cadherin was upregulated in breast cancer cells that normally express low E-cadherin and are highly invasive ^17^, whereas alignment decreased when E-cadherin was downregulated in cohesive epithelial cells. Moreover, E-cadherin expression in mouse keratinocytes was required to localize traction forces along the periphery of a cell aggregate; its downregulation redistributed traction to individual cells ^51^. Motivated by the idea that intercellular mechanical coupling can generate global matrix alignment and allow cells to migrate through a pre-strained matrix, we incorporated cell– cell adhesion into the model by setting *κ*_a,int_ = 10^−5^ N/m in Eq. 8 (Fig. S2). With cell–cell adhesion included, cell migration occurred preferentially in regions where the matrix had been locally strained (Fig. 4A, second from left labeled with “w/ adh, w/ *F*_tot,F_” and Movie S4). Matrix strain reached ∼ 0.5 and was maintained until ∼550 h (Figs. 4B, C). Intercellular adhesion increased both *F*_r_ and the radial bias of tensile forces (*F*_r_/*F*_θ_) throughout the simulation (Figs. 4D and S6B). This occurred because adhesion slowed cell escape from the cluster (Fig. 4E), allowing cells to collectively stiffen the matrix before migrating. As a result, cells experienced stronger durotactic cues in the radial direction, and 82% migrated with the radial migration bias greater than 0.8 (Fig. 4F).

In summary, our model suggests that cell–cell adhesion tunes migration paths in fibrous matrices (Fig. 5). Contractility from a single cell is not enough to produce anisotropic stiffening over matrix. When adhesion is weak across multiple cells, cells migrate immediately after seeding with greater directionality but matrix deformation is not persistent. When adhesion is strong, cells remain mechanically coupled until matrix stiffening generates sufficiently strong durotactic cues to overcome intercellular adhesion, enabling directed migration along the stiffened paths.

**Figure 5.**
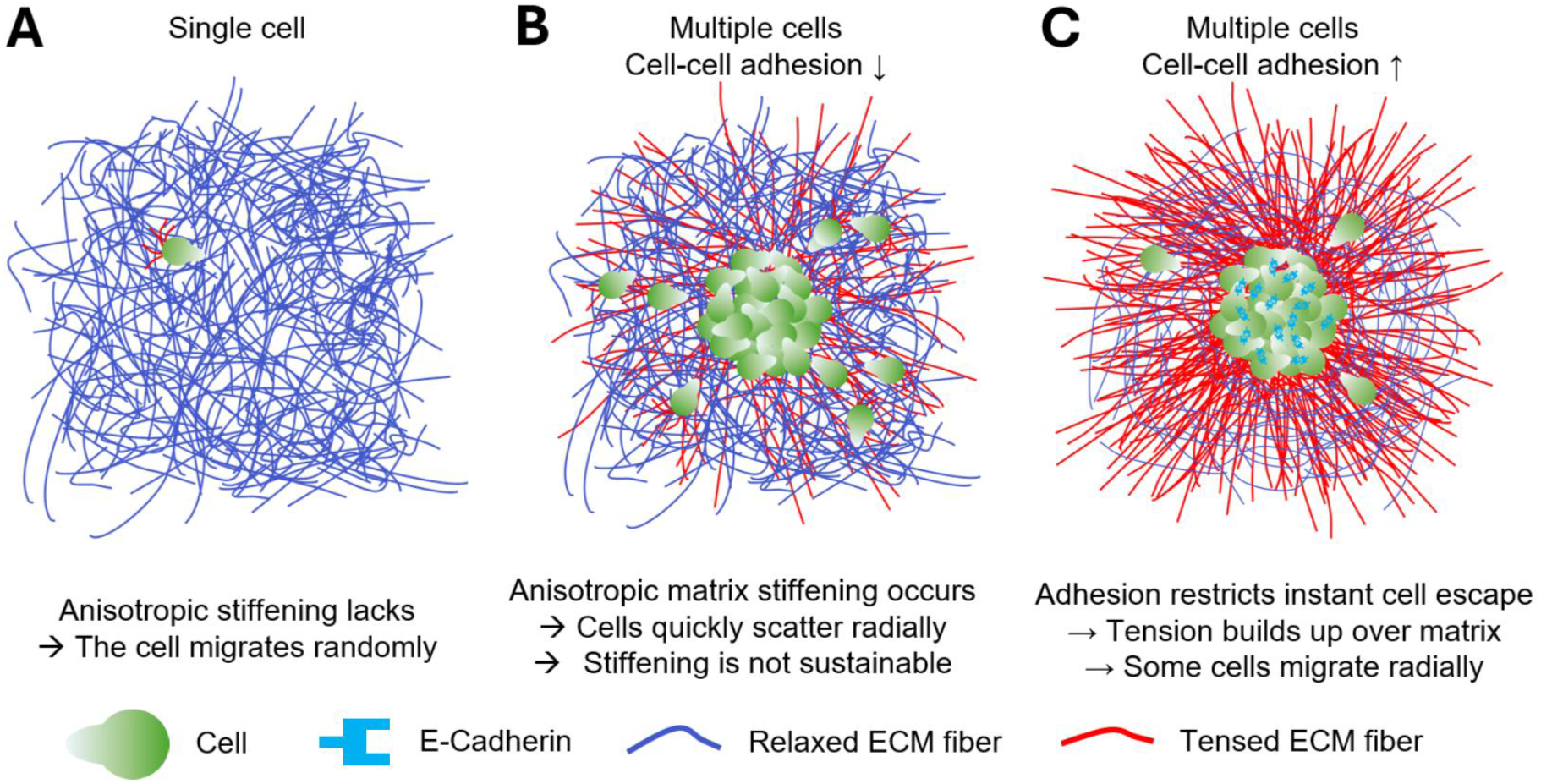
How cell migration can be enhanced by anisotropic stiffening and mechanical feedback. (A) When only a single cell is present, migration is not biased in any direction because forces generated by one cell are not high enough to induce anisotropic stiffening. (B) A cell cluster with no or weak cell-cell adhesion can anisotropically stiffen a matrix, resulting in radially biased migration. However, cells scatter quickly, and force level developed by the cells is not very high and not sustainable for a long time. (C) A cell cluster with strong cell-cell adhesion contracts a matrix very strongly, significantly enhancing radially biased migration. In addition, cells leave the cluster much slower, reminiscent of cancer cell migration during intravasation.

### Increased cell contractility promotes collective migration along migration paths

As with cell–cell adhesion, cell contractility is another critical determinant of fiber alignment around cell clusters and of subsequent migratory behaviors. Prior studies have shown that inhibition of Rho or ROCK suppresses collagen contraction and alignment in epithelial tissues ^9,17^ and tumor spheroids ^18^. Downregulation of actomyosin contractility also reduces cell spreading from clusters ^9^.

To examine how contractility influences matrix remodeling and migration in our model, we increased the contraction forces of both forward and rear cell-points by 50%. We observed clear formation of migration paths, indicating that cells with greater contractility escaped the cluster more frequently and were able to recognize paths previously stiffened by leader cells (Fig. 4A, third from left labeled with “w/ 1.5*F*_c_” and Movies S5, 6). This collective behavior requires frequent cell escape; otherwise, the matrix relaxes before follower cells arrive, causing the migration path to disappear. Indeed, tumor-cell studies have shown that leader cells align collagen fibers and generate cell-scale tracks that follower cells subsequently use for migration ^52,53^, consistent with earlier computational work demonstrating that a cell can follow the path established by a cell that began migrating earlier ^27^. The increased contractility strengthened mechanical coupling between cells and enabled 83% of cells to migrate with the radial migration bias greater than 0.8 when cell contractility increased (Fig. 4F). Notably, increased contractility alone (w/o adh, w/ 1.5 *F*_tot,F_) improved migration directionality to a level comparable to that achieved by intercellular adhesion alone (w/ adh, w/ *F*_tot,F_). Increased contractility alone did not enhance matrix contraction, increase *F*_r_/*F*_θ_, or delay cell escape (Figs. 4B–E and S6C, D) relative to the baseline-contractility condition (w/o adh, w/ *F*_tot,F_). Instead, the improvement in radial migration arose from a substantial increase in the fraction of cells entering the migration path. This indicates that higher contractility enhances the ability of follower cells to detect and migrate onto the stiffened tracks generated by leader cells, thereby promoting orchestrated migration.

However, the contributions of contractility and adhesion to directional migration were dependent, as evidenced by their non-additive effects on radial directionality (Fig. 4F). Together, these results suggest that intercellular adhesion and cell contractility regulate migratory trajectories through distinct yet shared mechanisms. Intercellular adhesion promotes coordinated matrix contraction, whereas increased contractility enhances follower recruitment onto pre-stiffened migration paths.

Lastly, we repeated the quantitative analysis along the circumference of circles with *R*_a_ = 300 µm (Figs. S7A, B) and 400 µm (Figs. S7C, D), instead of 200 µm from which previous conclusions were drawn (Figs. 4C, D), to confirm that our conclusions were not dependent on the selected circle size. Overall, at higher *R*_a_, matrix contraction stain was suppressed under all conditions, but the inclusion of cell-cell adhesion consistently better sustained matrix contraction for a longer time. Moreover, *F*_r_/*F*_θ_ tended to be higher when cell-cell adhesion was included. These observations were consistent regardless of the circle size used for the analysis.

## DISCUSSION

Cells continuously sense and respond to the mechanical properties of their microenvironment while they migrate. Durotaxis—directed cell migration toward regions with higher stiffness—has been widely observed in mesenchymal, epithelial, and cancer cell types. Durotaxis is known to play critical roles in wound healing, development, and tumor invasion ^54,55^. To reproduce the durotactic behavior of cells, various computational models have been developed ^4,5^. We have also introduced a durotaxis model that accounts for the viscoelastic nature of extracellular environments ^37^. Using our previous model, we demonstrated how the durotactic migration of a single cell is regulated by biophysical properties of the substrate, including elasticity, viscosity, and stiffness profile.

In this study, we examined the influence of anisotropic matrix stiffening on the durotactic migration of multicellular assemblies. To induce anisotropic stiffening, we first applied normal strain to the matrix in one direction, following prior experimental approaches that utilized prestrained matrices to investigate durotaxis ^44,46,49^. To characterize the mechanical properties of the matrix after strain application, we quantified the ratio of stiffness measured parallel to the applied strain to that measured in the perpendicular direction, as well as the extent of permanent deformation after unloading (Fig. 2) ^47,48^. We observed that cell migration exhibited a stronger bias toward the direction of applied strain as the stiffness ratio increased (Fig. 3). While this correlation is indicative of durotactic behavior, it is also possible that strain-induced fiber alignment contributed to the biased migration through contact guidance. However, further analysis demonstrated that the directed migration observed in our simulations was primarily driven by durotaxis rather than contact guidance.

Next, we probed cell-induced anisotropic stiffening of the matrix in the absence of externally applied strain (Fig. 4). When intercellular adhesion was absent, cells initially positioned at the center of the matrix migrated freely, producing only localized matrix deformations. In contrast to previous experimental observations ^17,18,33^, we did not observe large-scale matrix remodeling preceding cell migration under these conditions. When intercellular adhesion was incorporated, cells collectively contracted the matrix, resulting in the development of strong radial tensile forces. Both matrix contraction and tensile forces persisted for much longer durations in the presence of intercellular adhesion. Consequently, a subset of cells exhibited directed migration in the radially outward direction along stiffened matrix fibers. These cells could overcome intercellular adhesive forces by exerting traction on fibers with increased mechanical resistance. We interpret this behavior as durotactic migration mediated by mechanical feedback, wherein cells actively remodel and stiffen their extracellular environment and subsequently migrate preferentially along the stiffened region of the matrix.

Cells contract and remodel the extracellular matrix through actomyosin-generated contractile forces. Upregulation of cell–cell adhesion was reported to enhance ECM remodeling ^17,51^, whereas inhibition of myosin motor activity via perturbation of the Rho–ROCK signaling pathway was shown to suppress extracellular matrix contraction and fiber alignment ^9,17,18^. In our simulations, increasing cellular contractility enabled cells to remodel a matrix more substantially and overcome intercellular adhesion more readily, thereby increasing the fraction of cells migrating along stiffened matrix fibers. These migrating cells aligned along common trajectories, resembling the leader–follower organization reported in previous studies. Collectively, these results suggest that cell–cell adhesion and cell contractility regulate migratory trajectories through distinct yet complementary mechanisms. Specifically, intercellular adhesion promotes coordinated matrix contraction, whereas higher cell contractility facilitates the ability of follower cells to enter and persist along established migration paths.

Admittedly, our model does not account for all biological mechanisms known to play an important role in cell migration. During epithelial–mesenchymal transition, epithelial cells progressively lose intercellular adhesion and acquire mesenchymal phenotypes characterized by increased motility and invasiveness ^56^. In our framework, cell–cell adhesion was introduced via a simplified mechanism to delay cell escape and to enable cells to remodel the ECM collectively. The strength of cell–cell adhesion in our model was held constant throughout the simulation. As a result, cell escape did not arise from adhesion weakening; instead, some cells were detached from the cluster only after the surrounding matrix had stiffened sufficiently to overcome adhesion to neighboring cells. Another limitation of our model is the absence of force-dependent maturation of focal adhesions. In living cells, focal adhesions grow in size and become stabilized under tensile loading, thereby strengthening cell–matrix coupling and enabling more effective force transmission on stiffer substrates ^57-59^. Our framework does not fully capture how cells preferentially stabilize adhesions in mechanically supportive regions of the ECM. In addition, our matrix model does not fully incorporate the viscoelastic features of biological ECMs although it can capture the non-linear stress-strain relationship of ECMs. Experimental and theoretical studies have suggested that stress relaxation and plastic deformation arise, at least in part, from force-dependent rupture and reformation of cross-linking points between matrix fibers ^48,60^. By contrast, connections between matrix nodes in our model are assumed to be permanent. This assumption likely suppresses stress relaxation and limits the extent of plastic deformation that the matrix can undergo, potentially leading to an overly elastic response compared with real ECMs.

## CONCLUSION

In this study, we developed a mechanical model to investigate the mechanisms underlying durotaxis on a fibrous matrix. We demonstrated that anisotropic stiffening—whether generated by externally applied strain or by cell-mediated matrix remodeling—can bias cell migration toward specific directions. We further showed that intercellular adhesion and cellular contractility play essential roles in coordinating collective matrix contraction, which in turn facilitates directional migration. In future work, we aim to extend this framework to more faithfully capture cancer metastasis, incorporating cell-state transitions (e.g, epithelial-mesenchymal transitions) and additional stromal cell types such as cancer-associated fibroblasts, which play critical roles in tumor invasion.

