## Supplementary Material for "Durotactic Migration Driven by Anisotropic Matrix Stiffening and Mechanical Feedback"

**SUPPLEMENTARY FIGURES**

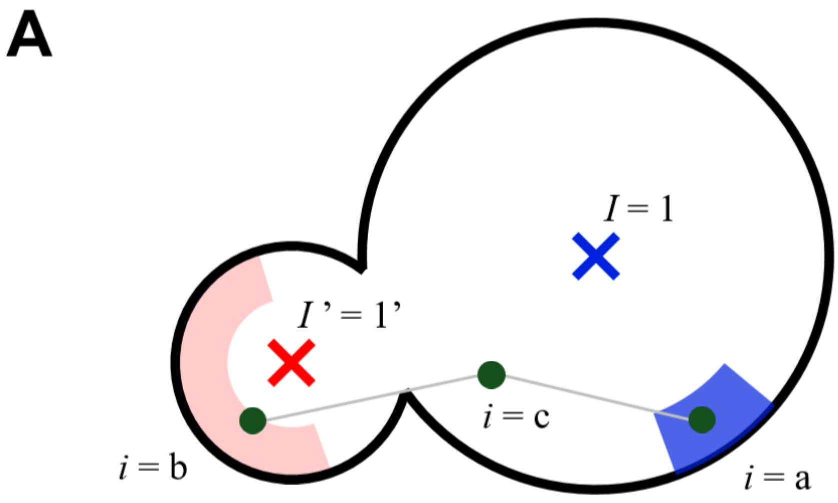


**B**

$$
\overset{\boldsymbol{M}}{\left[\begin{array}{cccc|cccccc}
\eta+\alpha+\lambda & -\alpha & 0 & 0 & -\lambda & 0 & 0 & 0 & 0 & 0 \\
-\alpha & \eta+\alpha+\lambda & 0 & 0 & 0 & -\lambda & 0 & 0 & 0 & 0 \\
0 & 0 & \eta+\alpha+\lambda & -\alpha & 0 & 0 & 0 & -\lambda & 0 & 0 \\
0 & 0 & -\alpha & \eta+\alpha+\lambda & 0 & 0 & 0 & 0 & -\lambda & 0 \\
\hline
-\lambda & 0 & 0 & 0 & \gamma+\xi+\lambda & 0 & -\xi & 0 & 0 & 0 \\
0 & -\lambda & 0 & 0 & 0 & \gamma+\xi+\lambda & -\xi & 0 & 0 & 0 \\
0 & 0 & 0 & 0 & -\xi & -\xi & \gamma+2\xi & 0 & 0 & 0 \\
0 & 0 & -\lambda & 0 & 0 & 0 & 0 & \gamma+\xi+\lambda & 0 & -\xi \\
0 & 0 & 0 & -\lambda & 0 & 0 & 0 & 0 & \gamma+\xi+\lambda & -\xi \\
0 & 0 & 0 & 0 & 0 & 0 & 0 & -\xi & -\xi & \gamma+2\xi
\end{array}\right]}
\overset{\boldsymbol{u}}{\left[\begin{array}{c}
v_{x,1} \\ v_{x,1'} \\ v_{y,1} \\ v_{y,1'} \\ \hline v_{x,a} \\ v_{x,b} \\ v_{x,c} \\ v_{y,a} \\ v_{y,b} \\ v_{y,c}
\end{array}\right]}
=
\overset{\boldsymbol{f}}{\left[\begin{array}{c}
F_{x,1} \\ F_{x,1'} \\ F_{y,1} \\ F_{y,1'} \\ \hline F_{x,a} \\ F_{x,b} \\ F_{x,c} \\ F_{y,a} \\ F_{y,b} \\ F_{y,c}
\end{array}\right]}
$$

**Figure S1. Example of the linear system formulation.** (A) A schematic showing a simple system with one cell comprising two cell-points ($I$ = 1 and $I'$ = 1’) and three matrix nodes ($i$ = a, b, and c). Two of the matrix nodes (a and b) lie within the activated adhesion sub-region of the two cell-points. By contrast, the other matrix node (*c*) does not interact with any of the cell-points but connects the matrix nodes a and b. (B) Linear system formulation in the form of $\boldsymbol{Mu} = \boldsymbol{f}$ (Eq. 2) for the simple system shown in (A). After $\boldsymbol{M}$ and $\boldsymbol{f}$ are updated at each time step, based on system configuration and force development, the velocity vector $\boldsymbol{u}$ is found by solving $\boldsymbol{M}^{-1}\boldsymbol{f}$ to update the location of each position.

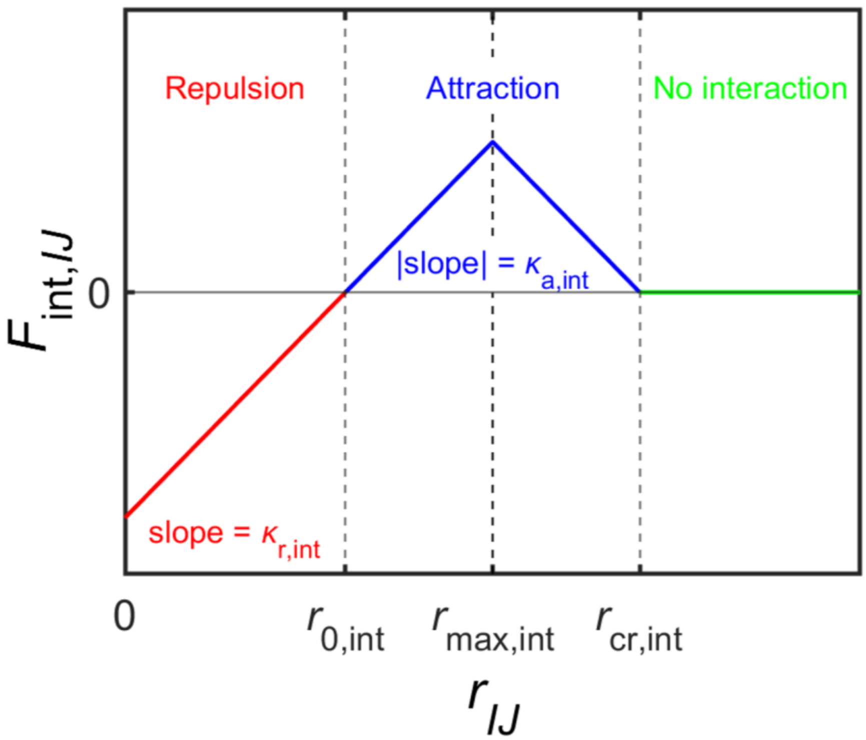


**Figure S2. Force acting between neighboring cells.** Cell-cell interaction force ($F_{\mathrm{int},IJ}$) is dependent on a cell-cell distance ($r_{IJ}$) (Eq. (8)). $r_{IJ}$ is divided into three regimes: i) repulsion when $r_{IJ}$ is shorter than a rest distance ($r_{0,\mathrm{int}}$), ii) attraction when $r_{IJ}$ lies between $r_{0,\mathrm{int}}$ and a critical distance ($r_{\mathrm{cr,int}}$), and iii) no interaction when $r_{IJ}$ is greater than $r_{\mathrm{cr,int}}$. In each regime, $F_{\mathrm{int},IJ}$ is then defined as a linearly increasing or decreasing function with repulsion strength ($\kappa_{\mathrm{r,int}}$) or attraction strength ($\kappa_{\mathrm{a,int}}$). Note that the sign of $\kappa_{\mathrm{a,int}}$ changes at $r_{\mathrm{max,int}}$ at which an attractive force becomes maximal.

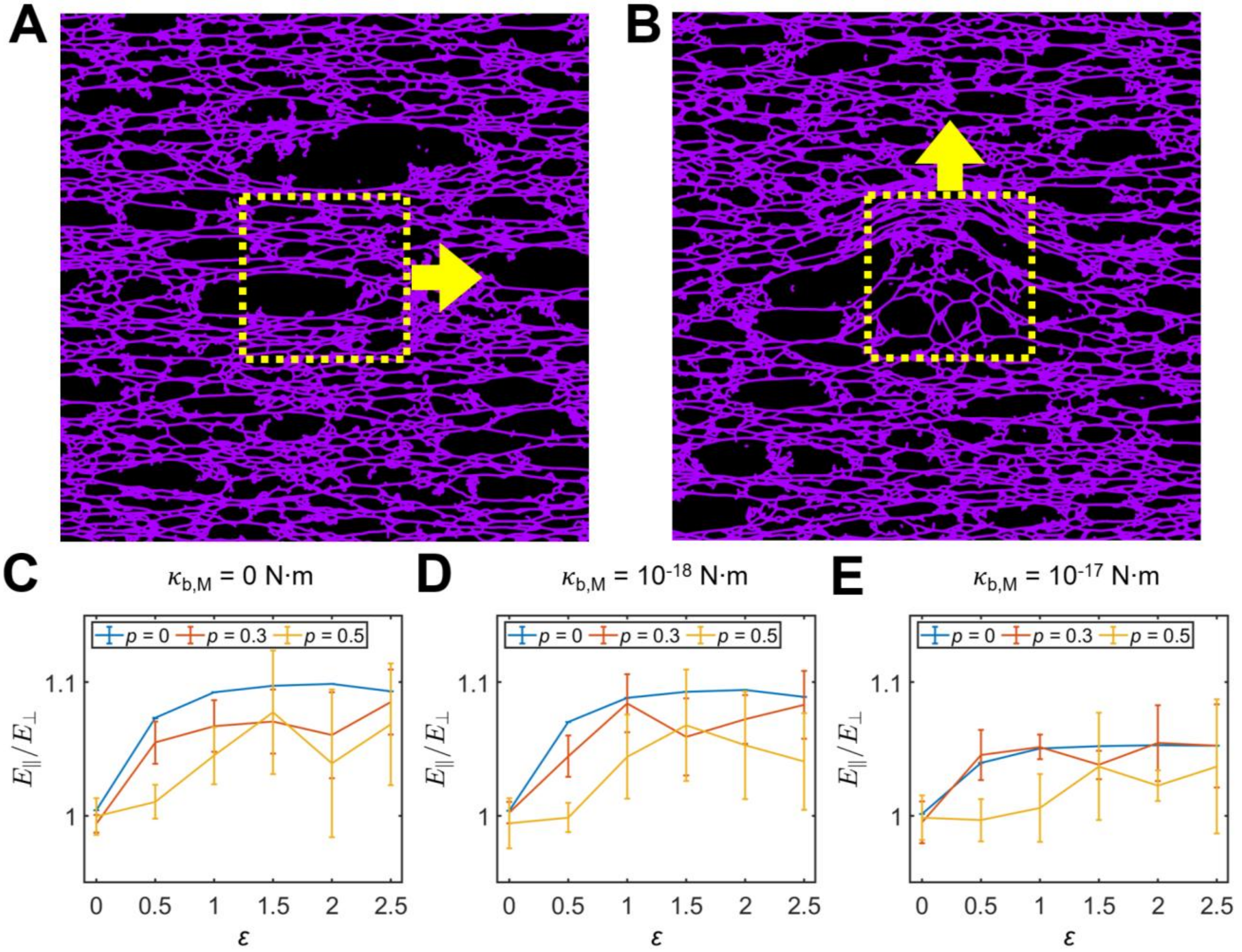


**Figure S3. Measurement of anisotropic stiffness for externally strained matrix.** (A,B) Matrix nodes located within a square (yellow dashed boxes) at the center of matrices are identified. An external force ($F_{\text{ext}}$) is applied to these matrix nodes in either (A) parallel or (B) perpendicular direction to the direction of externally applied strain, to measure stiffness in those directions ($E_{\parallel}$ and $E_{\perp}$). (A, B) In this example, $p$ = 0.5, $\varepsilon$ = 2, and $\kappa_{\text{b,M}}$ = $10^{-17}$ N·m. (C-E) $E_{\parallel}/E_{\perp}$ values were calculated with different values of $p$ and $\kappa_{\text{b,M}}$.

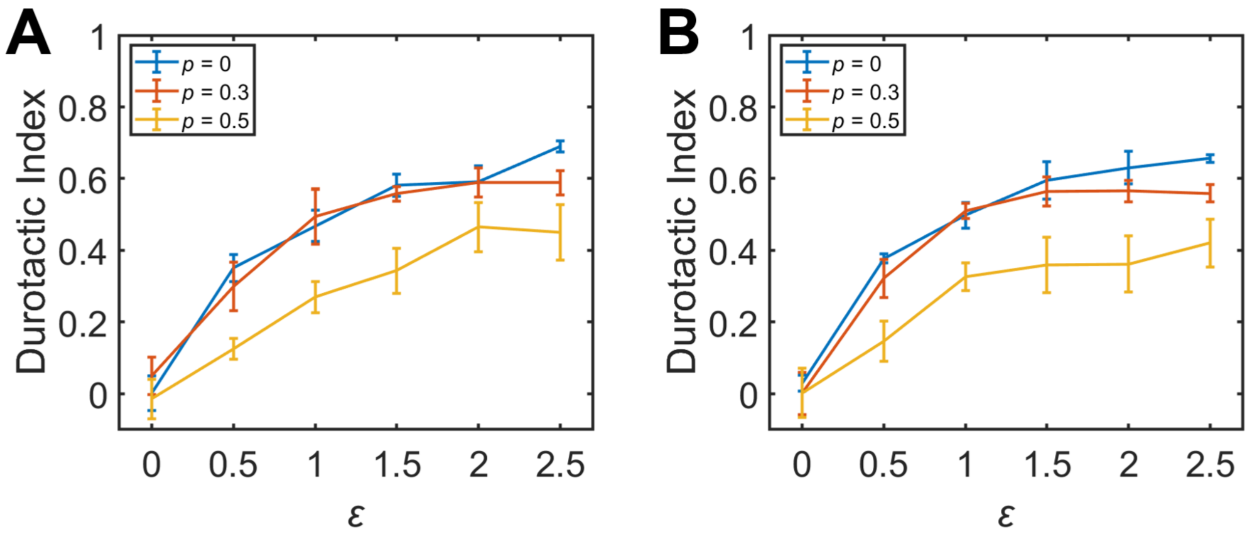


**Figure S4. Durotactic index measured with different bending stiffness ($\kappa_{b,M}$).** Durotactic index shown in Fig. 3C was measured with $\kappa_{b,M} = 10^{-17}$ N·m. (A) $\kappa_{b,M} = 0$ N·m. (B) $\kappa_{b,M} = 10^{-18}$ N·m.

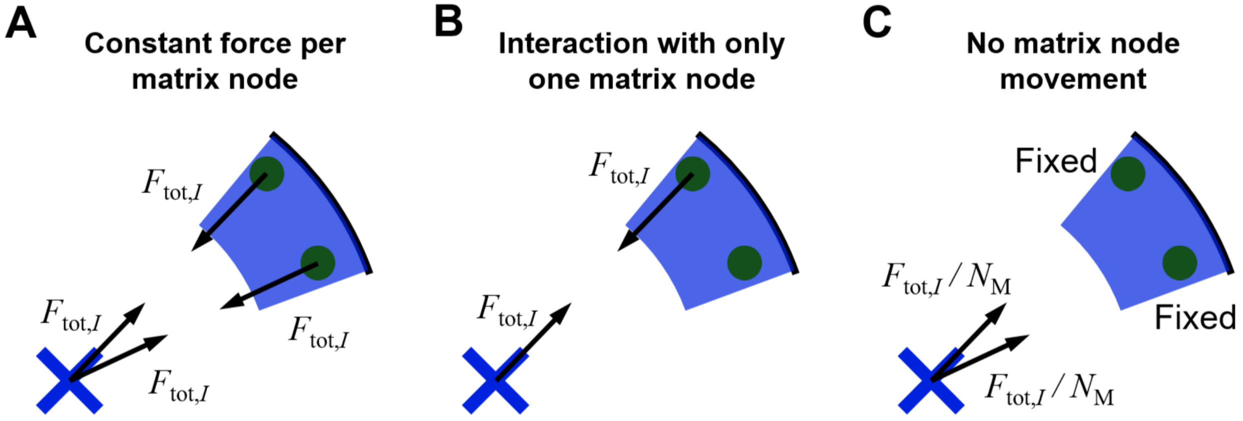


**Figure S5. Different modes of cell-matrix interaction.** (A) Cell-points exert a constant force to each matrix node, not the total force divided by the number of interacting matrix nodes. (B) The total contractile force is applied to a randomly selected single node within the lamellipodium-like adhesion region. (C) Matrix nodes are not allowed to move at all by contractile forces.

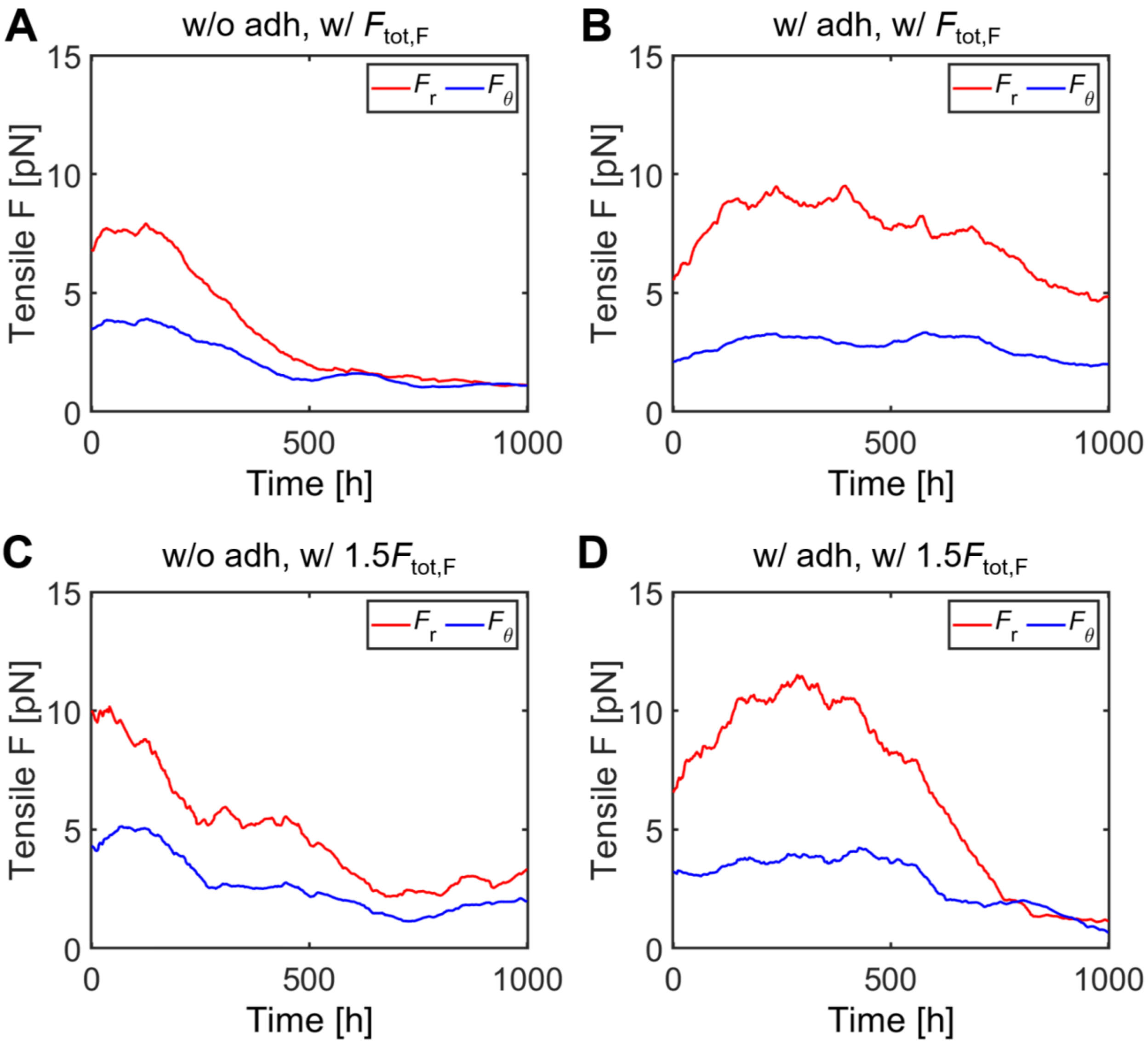


**Figure S6. Analysis of radial ($F_r$) and circumferential ($F_\theta$) tensile forces developed in circular matrices by cells.** $F_r$ and $F_\theta$ were measured over time under four different conditions: (A) w/o cell-cell adhesion (adh), w/ the reference total contractile force of the front cell-point ($F_{tot,F}$), (B) w/ adh, w/ $F_{tot,F}$, (C) w/o adh, w/ $1.5F_{tot,F}$, and (D) w/ adh, w/ $1.5F_{tot,F}$.

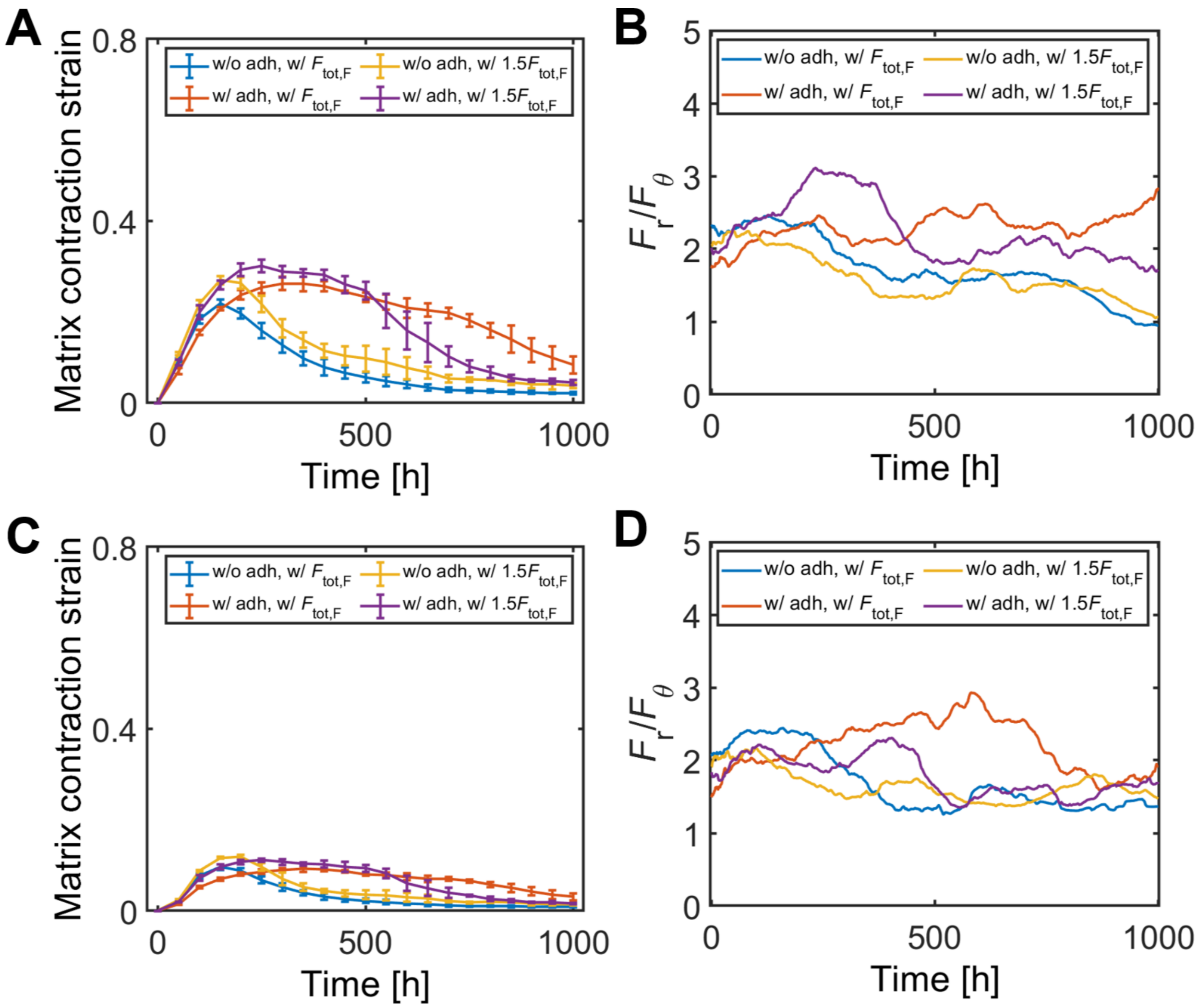


**Figure S7. Analysis of matrix deformation and forces measured with different circular region sizes ($R_a$).** (A, C) Matrix strain and (B, D) The ratio of radial forces to circumferential ones ($F_r/F_\theta$) were measured along the circumference of a circle centered at the domain center with (A, B) 300 μm or (C, D) 400 μm in radius.

### SUPPLEMENTARY TABLE

**Table. S1.** List of parameters employed in our model. Asterisk (*) indicates reference parameter values that were varied in some of simulations as noted in manuscript.

| Symbol | Definition | Value |
|---|---|---|
| $\eta_{\mathrm{F}}$ | Drag coefficient of front cell-points due to a medium | 0.036 [kg/s] |
| $\eta_{\mathrm{R}}$ | Drag coefficient of rear cell-points due to a medium | 0.036 [kg/s] |
| $\alpha$ | Drag coefficient between front and rear cell-points within each cell | 0.36 [kg/s] |
| $\beta$ | Drag coefficient between neighboring cell-points | 0.36 [kg/s] |
| $R_{\mathrm{out,F}}$ | Outer radius of front cell-points | $1.4\times10^{-5}$ [m] |
| $R_{\mathrm{in,F}}$ | Inner radius of front cell-points | $6.0\times10^{-6}$ [m] |
| $R_{\mathrm{out,R}}$ | Outer radius of rear cell-points | $1.0\times10^{-5}$ [m] |
| $R_{\mathrm{in,R}}$ | Inner radius of rear cell-points | $4.0\times10^{-6}$ [m] |
| $r_{0,\mathrm{FR}}$ | Equilibrium distance between front and rear cell-points | $8.0\times10^{-6}$ [m] |
| $\kappa_{\mathrm{s,FR}}$ | Extensional stiffness maintaining $r_{0,\mathrm{FR}}$ | 0.05 [N/m] |
| $\kappa_{\mathrm{r,int}}$ | Repulsive strength between cell-points | $1.0\times10^{-4}$ [N/m] |
| $\kappa_{\mathrm{a,int}}$ | Attractive strength between cell-points | 0 or $1.0\times10^{-5}$ [N/m] |
| $r_{0,\mathrm{int}}$ | Intercellular distance below which a repulsive force acts | Sum of $R_{\mathrm{in}}$'s |
| $r_{\mathrm{cr,int}}$ | Maximum intercellular distance for an attractive force | Sum of $R_{\mathrm{out}}$'s |
| $r_{\mathrm{max,int}}$ | Intercellular distance where an attractive force is maximized | $(r_{0,\mathrm{int}} + r_{\mathrm{cr,int}}) / 2$ |
| $\gamma$ | Drag coefficient of matrix nodes due to a medium | 0.0036 [kg/s] |
| $\xi$ | Drag coefficient between matrix nodes | 0.036 [kg/s] |
| $r_{0,\mathrm{M}}$ | Equilibrium length of matrix chains | $5.0\times10^{-6}$ [m] * |
| $\kappa_{\mathrm{s,M}}^{0}$ | Extensional stiffness of matrix chains | $1.0\times10^{-5}$ [N/m] |
| $\kappa_{\mathrm{b,M}}$ | Bending stiffness of a matrix | $1.0\times10^{-5}$ [N/m] * |
| $\theta_{0,\mathrm{M}}$ | Equilibrium angle between adjacent matrix chains | $\pi/3$, $2\pi/3$, or $\pi$ [rad] |
| $\lambda$ | Drag coefficient between cell-points and matrix nodes | 0.036 [kg/s] |
| $F_{\mathrm{tot,F}}$ | Total contractile force of front cell-points | $8.0\times10^{-10}$ [N] * |
| $F_{\mathrm{tot,R}}$ | Total contractile force of rear cell-points | $1.0\times10^{-10}$ [N] * |
| $p$ | Removal probability of matrix chains | 0.3 * |
| $\varepsilon$ | External strain level applied to matrices | 0, 0.5, 1, 1.5, 2, or 2.5 |
| $F_{\mathrm{ext}}$ | Magnitude of a force applied to selected matrix nodes for stiffness measurement | $1.0\times10^{-11}$ [N] |
| $\Delta t$ | Time step | 0.36 [s] |

### SUPPLEMENTARY MOVIES

**Movie S1. Cell migration on an unstrained matrix.** Cell trajectories (left, yellow) and individual cell locations (right, green) are separately shown on a matrix (purple) with $\varepsilon = 0$.

**Movie S2. Cell migration on an externally strained matrix.** Cell trajectories (left, yellow) and individual cell locations (right, green) are visualized separately on a matrix (purple) with $\varepsilon = 2$.

**Movie S3. Cell migration without cell-cell adhesion with the reference strength of contractile forces ($F_{tot,F}$).** Individual cells are visualized in green, and tensile forces developed on a matrix are visualized via color scaling.

**Movie S4. Cell migration with cell-cell adhesion and the reference strength of contractile forces ($F_{tot,F}$).** Individual cells are shown in green, and tensile forces acting on a matrix are visualized using color scaling.

**Movie S5. Cell migration without cell-cell adhesion with stronger contractile forces ($1.5F_{tot,F}$).** Individual cells are visualized in green, and tensile forces exerted on a matrix are indicated by color scaling.

**Movie S6. Cell migration with cell-cell adhesion and stronger contractile forces ($1.5F_{tot,F}$).** Individual cells are shown in green, and tensile forces developed on a matrix are shown via color scaling.
